# TRPM3 as novel target to alleviate acute oxaliplatin-induced peripheral neuropathic pain

**DOI:** 10.1101/2022.08.10.503165

**Authors:** Vincenzo Davide Aloi, Silvia Pinto, Rita van Bree, Katrien Luyten, Thomas Voets, Joris Vriens

## Abstract

Chemotherapy-induced peripheral neuropathic pain (CIPNP) is an adverse effect observed in up to 80% of cancer patients upon treatment with cytostatic drugs including paclitaxel and oxaliplatin. CIPNP can be so severe that it limits dose and choice of chemotherapy, and has a significant negative consequences on the quality of life of survivors. Current treatments for CIPNP are limited and unsatisfactory. TRPM3 is a Ca^2+-^permeable ion channel functionally expressed in peripheral sensory neurons involved in the detection of thermal stimuli. Here, we focus on the possible involvement of TRPM3 in acute oxaliplatin-induced mechanical allodynia and cold hypersensitivity. In vitro calcium microfluorimetry and whole-cell patch-clamp experiments showed that TRPM3 is functionally upregulated in both heterologous and homologous expression systems after acute (24h) oxaliplatin treatment, while direct application of oxaliplatin was without effect. In vivo behavioral studies using an acute oxaliplatin model for CIPNP showed the development of cold and mechano-hypersensitivity in control mice, which was lacking in TRPM3 deficient mice. In addition, the levels of pERK, a marker for neuronal activity, were significantly reduced in DRG neurons derived from TRPM3 deficient mice compared to control after oxaliplatin administration. Moreover, intraperitoneal injection of a TRPM3 antagonist, isosakuranetin, effectively reduced the oxaliplatin-induced pain behavior in response to cold and mechanical stimulation in mice with an acute form of oxaliplatin-induced peripheral neuropathy. In summary, TRPM3 thus represents a potential new target for the treatment of neuropathic pain in patients undergoing chemotherapy.

## 1. INTRODUCTION

Antineoplastic chemotherapeutic regimes have proven tremendously efficient in the regression of previously untreatable cancers. Chemotherapy, however, is highly notorious for its toxic effects on the central nervous system (CNS) including the progressive deterioration of learning, memory, and other cognitive skills [24], [36]. In addition, peripheral neuropathy is a potentially chronic and in specific cases also acute adverse event that occurs secondary to the treatment with various cytostatic drugs such as platinum based compounds, taxanes and vinca alkaloids. Neuropathic pain is a characteristic feature of peripheral neuropathy secondary to chemotherapy, which may be experienced in the form of paresthesia, burning or tingling sensations in the peripheral extremities [2],[5]. Overall, CIPNP has a significant negative effect on the quality of life of cancer survivors. Platinum-based compounds such as oxaliplatin and cisplatin have the strongest correlation with peripheral neuropathy [1]. Whereas the majority of antineoplastic drugs lead to a delayed development of neuropathy, oxaliplatin and paclitaxel can induce an acute pain syndrome, which can become chronic after long-term treatment.

Several members of the transient receptor potential (TRP) superfamily of cation channels are functionally expressed at the sensory endings of dorsal root ganglia (DRG) neurons, where they have been implicated in the transmission of a diverse range of mechanical, chemical and thermal sensory stimuli. With the presence of a leaky barrier between the blood and DRGs, it has been speculated that chronic exposure to antineoplastic drugs, in particular platinum based compounds, could lead to a modified activity of TRP channels in sensory DRG neurons [27]. Especially, treatment with oxaliplatin triggers the upregulation of the nociceptors TRPV1, TRPA1 and TRPM8 in DRG neurons, which may contribute to oxaliplatin-induced thermal hypersensitivity [3]. Paclitaxel has been found to induce an increased release of Substance P and other neuropeptides responsible for inducing hyperalgesia in the peripheral extremities [28]. The current options to prevent or treat CIPNP are narrow and rather limited to duloxetine [23] and tramadol [17]. Based on the relative abundance of TRP channels identified in DRG, and their potential role in mediating CIPNP, TRP channels represent promising novel targets for treating chemotherapy-induced peripheral pain [12],[18]. Recently, TRPM3 was identified as a novel sensory TRP channel expressed at molecular and functional level in a large subset of C and Aδ sensory neurons of mouse [34] and human [32]. TRPM3 is a polymodally gated ion channel which can be activated by different chemical substances such us the endogenous neurosteroid pregnenolone sulphate (PS) and by heat [34]. Furthermore, TRPM3 activation is well described to induce pain and stimulates the release of neuropeptides like CGRP [7]. In addition, TRPM3 is already well described to be involved in different types of pain like inflammatory [34] neuropathic pain and spontaneous pain after nerve injury [26]. Hence, given the functional expression of TRPM3 in somatosensory neurons and its role as nociceptor channel, this study aims to clarify the potential role of TRPM3 in CIPNP and to investigate its potential as a target to alleviate chemotherapy-induced neuropathic pain.

## 2. MATERIALS AND METHODS

### 2.1. Cell culture and transfection

HEK293T cells stably expressing murine *Trpm3* (HEK-mTRPM3) and non-transfected HEK293 (NT) were designed and cultured as described previously [34]. HEK293T cells stably transfected with human TRPM3 were developed and validated by SB Drug Discovery (Glasgow, UK). The human TRPM3 was stably expressed in HEK293T cells (HEK-hTRPM3) after induction with Tetracycline (3μg/ml). For transient transfection, cells were transfected with 2 μg of DNA 24 - 48 hours before measurement using TransIT transfection reagents (Mirus). DRG neurons from adult (postnatal weeks 8–12) mice were isolated and cultured as described elsewhere [7].

### 2.2. Calcium imaging and electrophysiology

HEK-mTRPM3 cells or DRG neurons were loaded with Fura-2-acetoxymethyl for 30 min at 37°C. Fluorescence was measured during alternating illumination at 340 and 380 nm using Eclipse Ti (Nikon) fluorescence microscopy system, and absolute calcium concentration was calculated from the ratio of the fluorescence signals at these two wavelengths (R = F340/F380) as [Ca^2+^]=Km × (R-R_min_)/(R_max_-R), where Km, Rmin and Rmax were estimated from in vitro calibration experiments with known calcium concentrations [30]. The standard external solution contained (in mM) 150 NaCl, 10 Hepes, 2 CaCl2, and 1 MgCl2 (pH 7.4 with NaOH). Whole-cell patch clamp experiments were performed with an EPC-10 amplifier and PatchMasterPro Software (HEKA Elektronik, Lambrecht, Germany). The sampling rate for the current measurements was 20 kHz and currents were digitally filtered at 2.9 kHz. The extracellular solution contained (in mM), 150 NaCl, 1 MgCl2, and 10 Hepes (pH 7.4 with NaOH), and the standard internal solution contained (in mM) 100 CsAsp, 45 CsCl, 10 EGTA, 10 Hepes, and 1 MgCl2 (pH 7.2 with CsOH).

### 2.3. Sensory nerve conduction

Sensory nerve conduction studies were performed as described previously [29]. Along the dorsal caudal nerve, sensory nerve action potentials (SNAPs) were measured by using platinum coated intramuscular wire electrodes (Technomed Europe, Maastricht, The Netherlands) and a Natus UltraPro S100 (Natus Medical Incorporated, Pleasanton, USA).

### 2.4. RT–qPCR

Methods were based on established protocols from our research group [33]. RNA extraction of murine DRGs neurons was done using RNeasy mini kit (QIAGEN). cDNA was generated from 1 μg of RNA using the First-strand cDNA Synthesis Kit (Thermofisher Scientific). RT-qPCR was performed on triplicate cDNA samples using specific TaqMan gene expression assays in the StepOne PCR system. Data is represented as mean ± SEM of 2(-ΔCt) for which ΔCt = Ct_gene of interest_ – Ct_geometric mean of endogenous controls_.

### 2.5. Animals

For all experiments 8-12 week old male C57BL/6J (Janvier Labs, Le Genest-Saint-Isle, France) mice were used. To investigate the effects of TRPM3 deficiency, homozygous (*Trpm3^-/-^*) mice were generated as described previously [34]. Mice of all genotypes were housed under identical conditions, with a maximum of five animals per cage, on a 12-h light–dark cycle. Food and water were provided *ad libitum*.

### 2.6. Drugs

Oxaliplatin was purchased from (Tocris Bioscience) and freshly dissolved in 5% glucose solution. For the acute oxaliplatin model mice received a single intraperitoneal administration of oxaliplatin (6 mg/kg), or vehicle [4]. Tramadol hydrochloride (5mg/kg, Sigma) was prepared in a mixture of (1% Hydroxypropylmethylcellulose HPMC/ 0.5 % Tween 80) and administered via oral gavage. Isosakuranetin (2mg/kg) was dissolved in Miglyol 812 (Caesar & Loretz GmbH, Hilden, Germany) containing 0.1% DMSO, and compounds or the vehicle alone were injected intraperitoneally. Pregnenolone sulphate was purchased from (Sigma-Aldrich) and dissolved in DMSO. Stock solutions of Pregnenolone sulphate was 100 mM.

### 2.7. Ethical approval

All animal experiments were carried out in accordance with the European Union Community Council guidelines and approved by the local Ethics Committee of the KU Leuven (P108/2016).

### 2.8. Behavioral tests

#### 2.8.1. Chemogenic pain model

PS (250 μM) was dissolved in PBS + 0.1% DMSO. The mice were allowed to acclimate in a clear plastic box for at least 40 min after which 20 μl of PS solution (5 nmol) was injected intraplantarly using a 30G needle coupled to a Hamilton syringe, and behavior was recorded. Experiments were performed during the light cycle. The duration of the PS-evoked nocifensive behavior such as licking and flicking was quantified for 10 minutes [34].

#### 2.8.2. Acetone spray test

Cold sensitivity was tested by the acetone spray test, as previously described [10]. Briefly, thirty minutes prior to the start of the experiments, all the animals were placed in a testing apparatus consisting in eight chambers with a mesh floor in order to acclimatize the mice with the environment. After the acclimation a volume of (50μl) of acetone was applied to one of the hind paw and the responses were monitored for 60 seconds post application. Responses to a cold stimulation were graded according to the following 5-point scale: 0-no response; 1-brief lift, sniff, flick, or startle; 2-jumping, paw shaking; 3-multiple lifts, paw lick; 4-prolonged paw lifting, licking, shaking, or jumping; 5-paw guarding [10].

#### 2.8.3. electronic Von Frey Assay

Mechanical allodynia was assessed by von Frey test using an electronic device (Bioseb, France) as previously described [14],[20]. Mice were placed individually in an eight-place chamber with a mesh floor and acclimatized for at least 30 minutes prior testing. Each paw was poked 4 times with a non-hygroscopic polypropylene von Frey tip of uniform diameter (0.8 mm) and the paw withdrawal threshold was calculated as the average of these four consecutive measurements. The numeric value of the force required to induce the paw withdrawal was recorded automatically by the apparatus.

### 2.9. Immunohistochemistry

Immunohistochemical staining of p-ERK on DRG neurons was performed based on a previously described protocol [26]. Firstly, after euthanizing the animals via CO2 inhalation, DRG neurons of each distinct group were dissociated, followed by their post fixation in 4 % paraformaldehyde solution for 4 hours at 4°C, and treated overnight at 4°C with the primary antibody (rabbit anti-pERK; 1:200, PhosphoSolutions). The pre-treated sections were then incubated for 2 hours at room temperature with goat anti-rabbit conjugated to Cy3. Triple washing with PBS was performed between each step. Subsequently, the sections were fixed with 4’,6-diamidino-2-phenylindole (DAPI) mounting media (Sigma-Aldrich). Immunofluorescence-labeled cells were imaged using a fluorescence microscope Eclipse Ci (Nikon). The NIH ImageJ software was used to quantify the labeled cells. pERK positive cells were defined as DAPI positive and a pERK immunofluorescence intensity that was above the threshold value of two times the mean value of the five lowest immunofluorescent pERK signals.

### 2.10. Experimental design and statistical analysis

Sample sizes for patch clamp experiments on HEK293T cells were based on earlier work from our team and literature, with a minimal sample size of n = 5. Sample sizes for calcium imaging on HEK293T cells and DRG neurons was based on literature, but always included a minimum of 5 independent experiments containing >100 cells per experiment. The sample sizes for all the *in vivo* experiments was based on a power analysis using statistical data from pilot experiments, aimed at detecting a difference of at least 20% between groups with an α of 0.05 and a power of 0.8. Data analysis, statistics and data display were performed using Origin 9.1 (OriginLab). All data are presented as the mean ± SEM from n biological replicates. Normality was tested using the Shapiro-Wilk test. The specific statistical tests used to determine significance of the differences between experimental data sets are indicated in the figure legends. *p* values of <0.05 were considered significantly different.

## 3. RESULTS

### 3.1. Effect of oxaliplatin on TRPM3 channel activity

First, we tested whether oxaliplatin has a direct agonistic effect on TRPM3 activity using microfluorimetric calcium imaging in HEK-mTRPM3 and HEK-hTRPM3 cell lines (Supplementary Fig.1). Direct application of oxaliplatin (100 μM) did not evoke any response in the intracellular Ca^2+^ concentration, while stimulation with PS (40 μM) induced a strong TRPM3–mediated Ca^2+^ influx. Similar results were obtained in isolated DRG neurons from mice (Supplementary Fig.1). Next, we investigated the effect of long-term incubation with oxaliplatin (100 μM for a period of 24h) on TRPM3 expression and activity, in HEK-mTRPM3, HEK-hTRPM3 cell lines and in DRG sensory neurons. In transiently transfected HEK293 cells with murine TRPM3, we measured significantly increased basal calcium levels at 37°C in cells treated with oxaliplatin compared to vehicle treated cells (Fig. 1A), and these increased cytosolic calcium levels could be partially reversed by application of the TRPM3 inhibitor primidone (100 μM) (Fig. 1A, B) [11]. Moreover, the [Ca^2+^]_i_ responses to PS or heat were significantly larger in oxaliplatin pre-incubated cells compared to vehicle treated cells (Fig. 1C-F). Similar results were obtained in HEK-hTRPM3 (Supplementary Fig. 2). These observations were further validated via whole-cell patch-clamp experiments, showing larger PS-induced current amplitudes in oxaliplatin pretreated HEK-mTRPM3 cells compared to vehicle controls (Fig. 2).

**Fig. 1:**
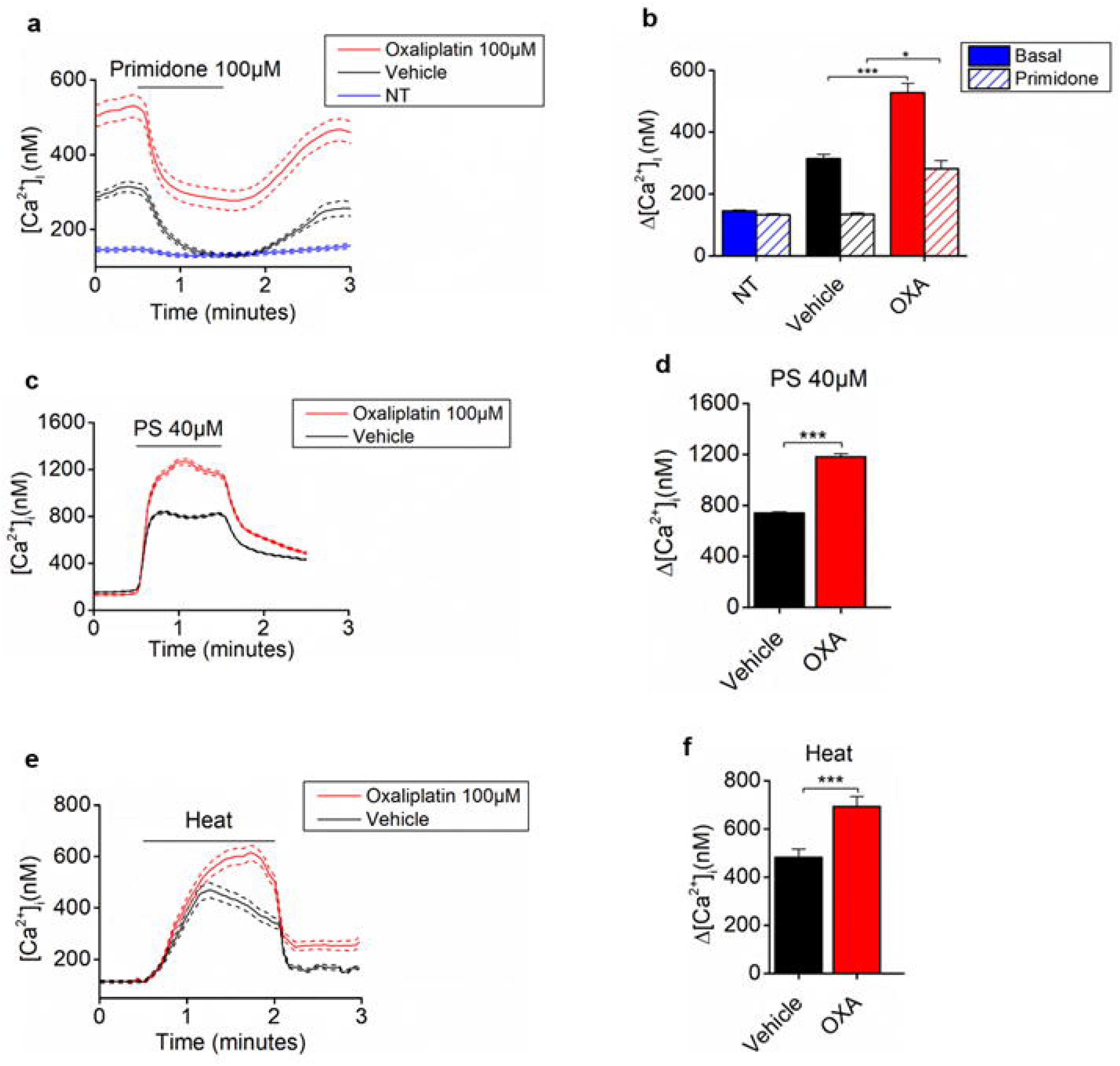
Modulation of TRPM3 function in HEK293 cells stably expressing murine TRPM3 after oxaliplatin pretreatment. (**A**) Time course of intracellular calcium concentrations ([Ca^2+^]_i_) (mean ± SEM) at 37°C upon application of the TRPM3 inhibitors primidone (100 μM) in HEK293-mTRPM3T cells after pretreatment with oxaliplatin (100μM) (n=299) and vehicle (n=293), and non-transfected (NT) cells (n = 97) (N = 3 independent experiments). (**B**) Basal intracellular calcium concentrations before (full bars) and after application of primidone (open bars). Data are represented as mean ± SEM. Statistically significant changes in basal channel activity were assessed using a Kruskal-Wallis ANOVA with Dunn’s posthoc test, where *p=<0.05 and ***<p=0.001. (**C** and **E**) Time course of intracellular calcium concentrations ([Ca^2+^]_i_) (mean ± SEM) for HEK-mTRPM3 cells in response to PS (40μM) and heat (37°) after oxaliplatin (100 μM) and vehicle treatment. (**D** and **F**) Quantification of calcium responses for experiments as in panel C and E respectively. Where ***p<0.001 (Mann-Whitney t test).

**Fig. 2:**
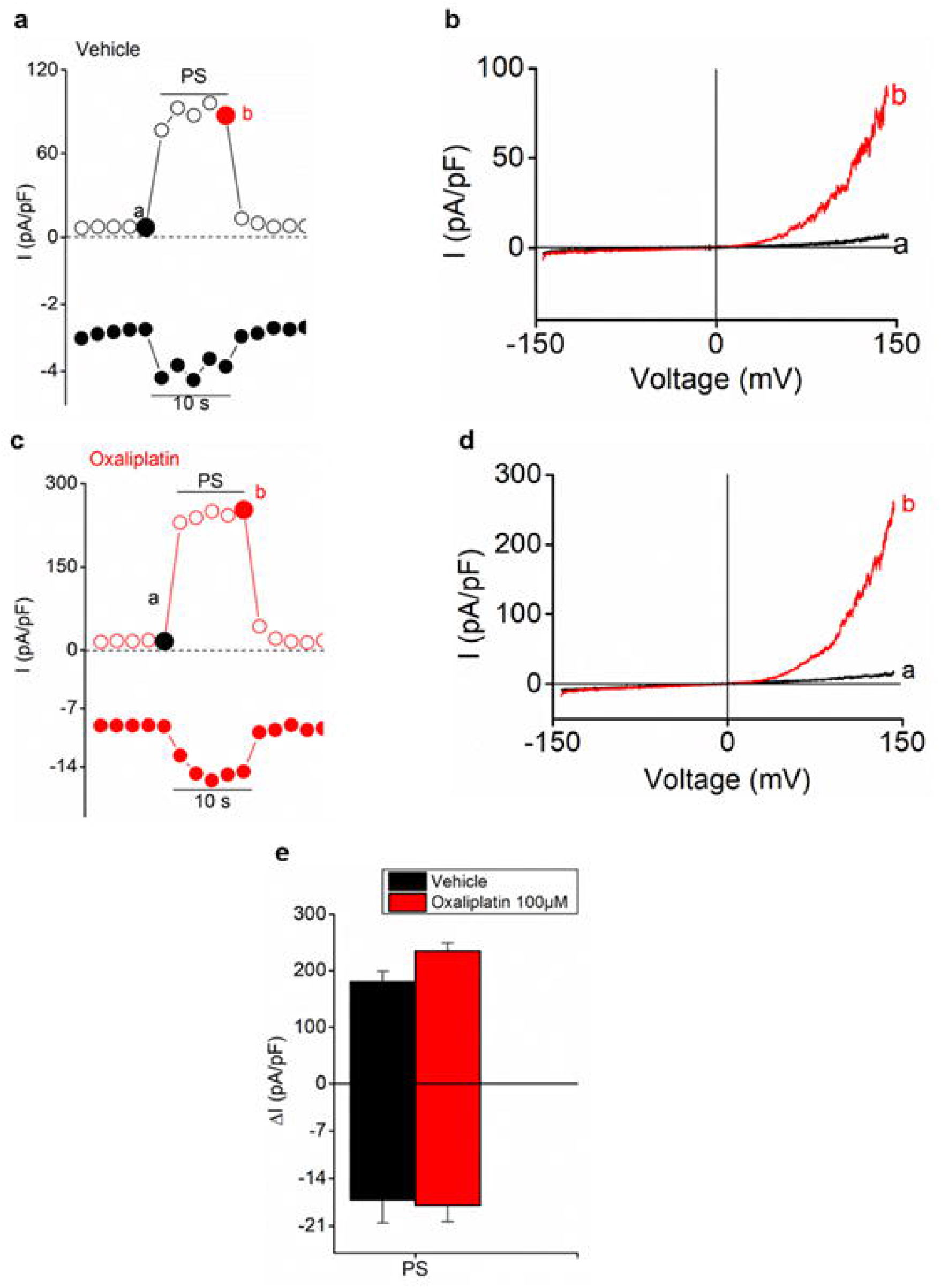
Modulation of TRPM3 function in HEK293-mTRPM3. (**A**) Time course at ± 150 mV of a whole-cell patch-clamp recording showing the effect of pregnenolone sulphate (PS; 40μM) on HEK-mTRPM3 expressing cells after vehicle pre-incubation (n = 16). (**B**) Current (I) – Voltage (V) relationship corresponding to the time points indicated in A. (**C**) Time course at ± 150 mV of a wholecell patch-clamp recording showing the effect of PS (40 μM) on HEK-mTRPM3 expressing cells after 24hr of oxaliplatin (100 μM) pre-incubation (n=16). (**D**) I-V relationship corresponding to the time points indicated in C. (**E**) Mean current increase at ± 150 mV in HEK-TRPM3 cells upon PS application after 24hr of vehicle or oxaliplatin pre-incubation. Where *p<0.05 (Mann-Whitney t test).

We also tested the effect of oxaliplatin pre-treatment on the PS responses in primary cultures of DRG neurons derived from wild type mice. Application of PS (40μM) to neurons evoked a Ca^2+^ response that was significantly larger in oxaliplatin-pretreated cells compared to vehicle controls (Fig. 3A, B). Moreover, the percentage of PS-responsive DRG neurons was increased after oxaliplatin treatment compared to vehicle controls (Fig. 3C). Interestingly, similar results were obtained when comparing DRG neurons isolated from mice at day 1 after oxaliplatin treatment (6 mg/kg) compared to vehicle (5% glucose) treated mice (Fig. 3D-F). Note that this *in vivo* treatment of the mice with a single dose of oxaliplatin had no effect on electrophysiological parameters of sensory nerve conduction (Supplementary Fig. 3), indicating that the integrity of the sensory nerves was preserved.

**Fig. 3:**
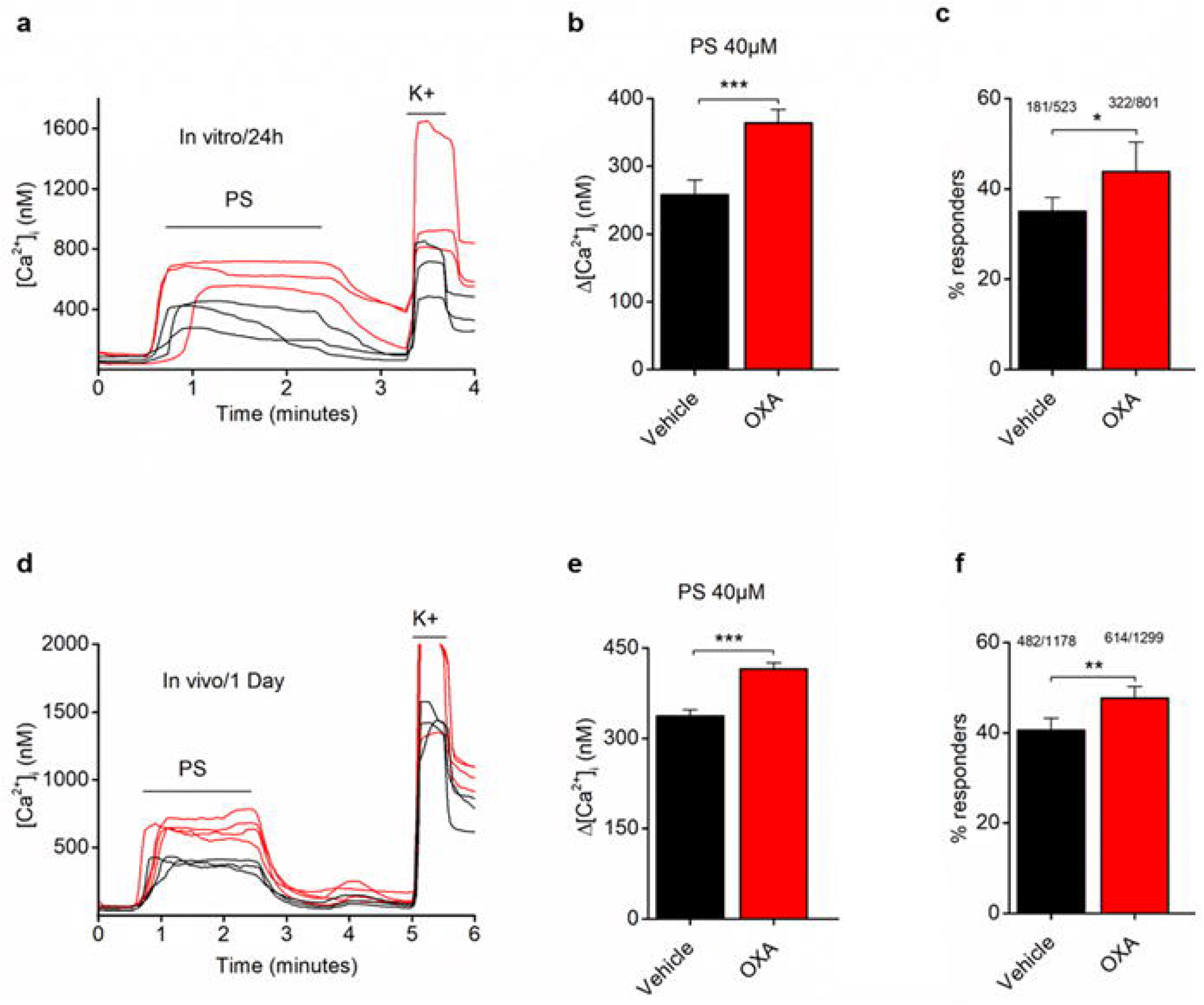
Modulation of TRPM3 function in DRG sensory neurons after oxaliplatin pretreatment. Time course of the intracellular Ca^2+^ concentration ([Ca^2+^]_i_) (mean ± SEM) in DRG isolated from wildtype animals TRPM3^+/+^ in response to PS (40 μM) after oxaliplatin (100μM; red) and vehicle treatment in black (**A**) and after 24h post oxaliplatin (6mg/kg; red color) or vehicle i.p injection (black color) (**D**). (**B** and **E**) Quantification of calcium responses for experiments as in panel A and B respectively, ***P<0,001 (Mann-Whitney t test). Percentage of sensory neurons derived from TRPM3^+/+^ responding stimulation by PS (40 μM) after oxaliplatin and vehicle pre-incubation (**C**) and after 24h post oxaliplatin (6mg/kg) or vehicle i.p injection (**F**). Where *p<0.05 (Chi-square test); ** p<0.01 (Chisquare test).

In order to elucidate whether the increased TRPM3 functionality was a result of increased mRNA expression, RT-qPCR was performed on DRG sensory neurons isolated from mice 24h after oxaliplatin or vehicle injection. This analysis did not reveal any significant change in TRPM3-encoding mRNA, but confirmed increased mRNA expression for TRPM8, in line with earlier work [9]. A similar outcome was also observed on HEK-mTRPM3 after 24hr of oxaliplatin pre-incubation. (Supplementary Fig. 4).

### 3.2. Oxaliplatin modulates TRPM3-dependent pain responses in vivo

In order to investigate whether oxaliplatin-induced peripheral neuropathic pain is associated with alterations in TRPM3-dependent pain responses, we compared the *in vivo* TRPM3 function in oxaliplatin-treated and vehicle-treated mice. To this aim, animals were submitted to a TRPM3-specific chemogenic pain model consisting of the intraplantar injection of PS (20 μl, 5nmol) [34]. The nociceptive response was quantified as the total time of nociception, i.e. the time during which the animals licked, lifted or guarded the affected paw. Before oxaliplatin injection, the intraplantar injection of PS evoked robust nocifensive behavior in wild type mice, which was not observed in TRPM3-deficient mice, similar to earlier studies [34]. Interestingly, the PS-induced pain response in wild type mice was significantly enhanced at day 6 post injection with oxaliplatin (6mg/kg) compared to vehicle treated mice or to the pre-treatment condition. Even after oxaliplatin treatment, TRPM3-deficient mice remained non-responsive to PS, confirming the TRPM3-specificity of the assay (Fig. 4). Taken together, these data indicate that TRPM3 is functionally upregulated in DRG neurons after oxaliplatin treatment.

**Fig. 4:**
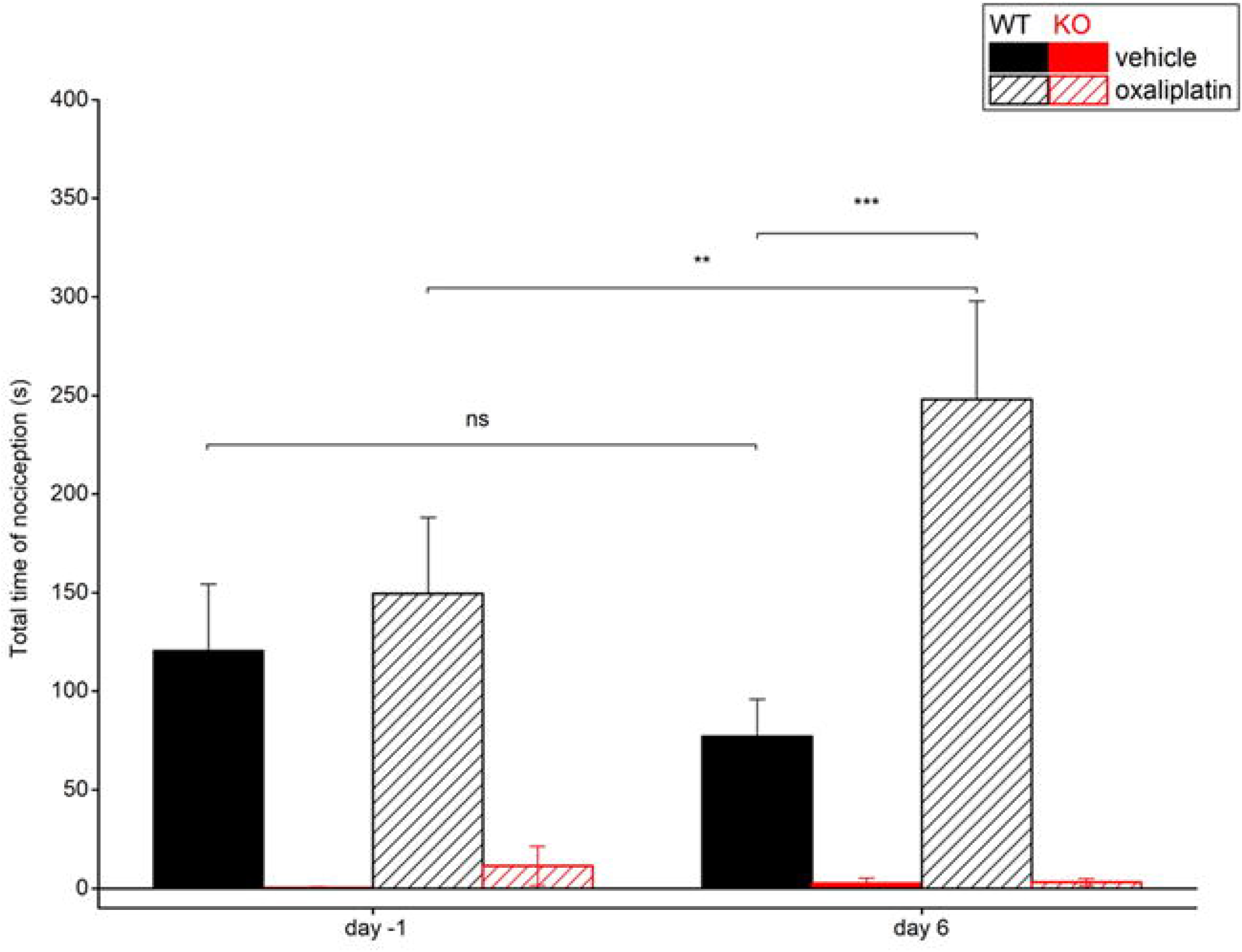
Nocifensive Responses to PS after oxaliplatin and vehicle injection. (**A**) Total duration of nocifensive behavior (paw licks and lifts in 2 min) in response to intraplantar injection of pregnenolone sulfate (PS, 5 nmol/paw) in TRPM3^+/+^ and TRPM3^-/-^ mice (n = 8 for each genotype). Data are represented as mean ± SEM. Statistically significant changes in the duration of PS-evoked pain responses were assesed using the Two-way ANOVA repeated measurement statistical test with Sidak-holm posthoc test. Where ** p<0.01; ***p<0.001

### 3.3. Involvement of TRPM3 in oxaliplatin-induced mechanical allodynia and cold hypersensitivity

To evaluate whether TRPM3 is involved in oxaliplatin-induced cold hypersensitivity and mechanical allodynia, we tested the effect of a single administration of oxaliplatin on behavioral responses to mechanical and cold stimuli in control and TRPM3^-/-^ mice. In wild type animals, the withdrawal threshold to a mechanical stimulus was significantly reduced at day 1 and day 6 after oxaliplatin treatment compared to the pre-treatment levels (Fig. 5A). Notably, oxaliplatin-induced mechanical hypersensitivity was strongly suppressed in TRPM3^-/-^ mice (Fig. 5A). Similarly, wild type mice exhibited significant and robust cold hypersensitivity in the acetone spray test at day 1 and day 6 following oxaliplatin treatment, whereas TRPM3^-/-^ mice did not develop such cold hypersensitivity (Fig. 5B).

**Fig. 5:**
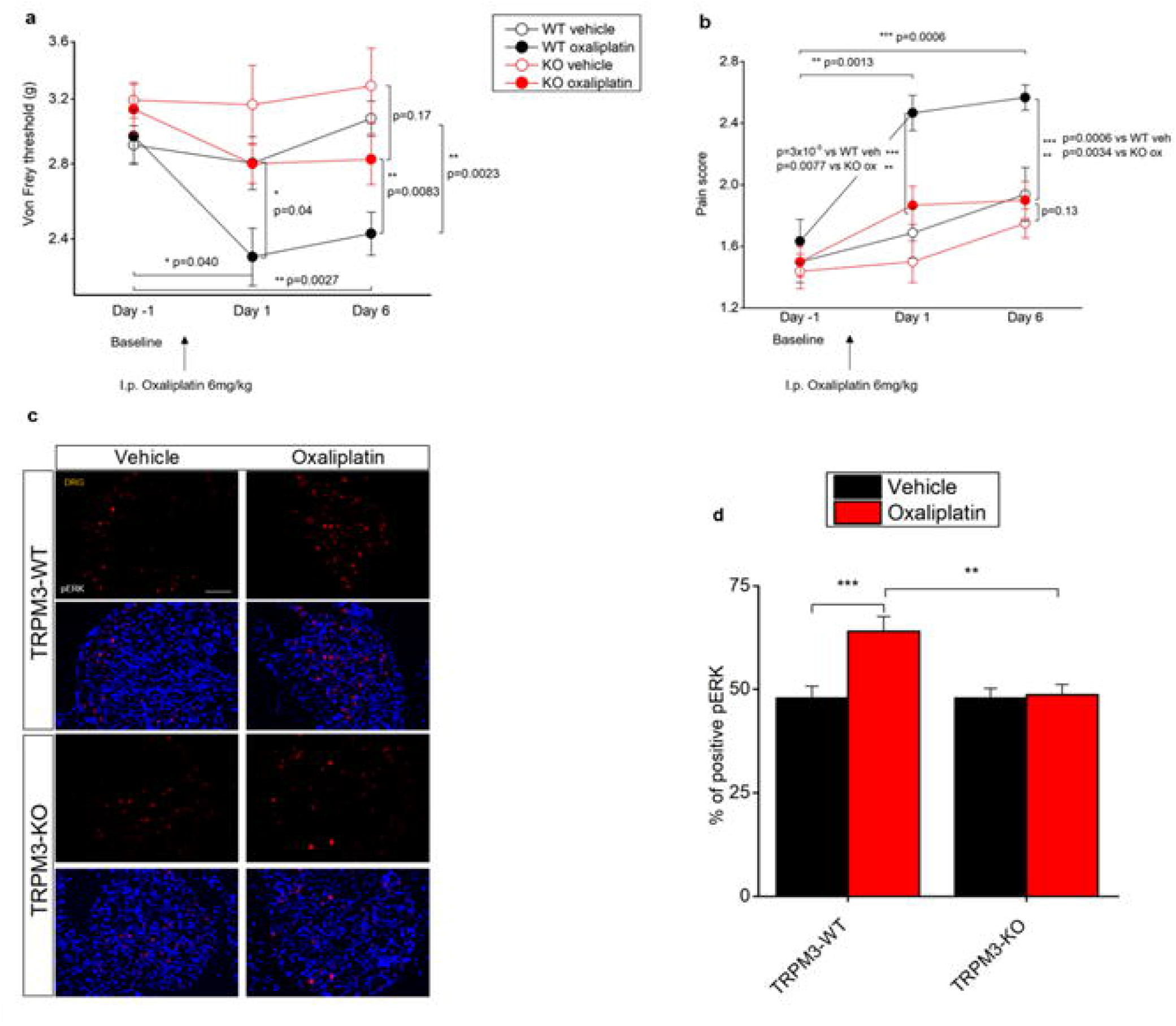
TRPM3 genetic ablation reduces oxaliplatin-induced mechanical allodynia and cold hypersensitivity and effects pERK expression in DRG neurons. (**A, B**) A single intraperitoneal injection of oxaliplatin (6mg/kg) induces in TRPM3^+/+^ mice (black color) a time-dependent reduction in mechanical nociceptive threshold, and increase in cold sensitivity respectively. The development of mechanical and cold allodynia observed in TRPM3^+/+^ animals after oxaliplatin treatment is decreased in TRPM3^-/-^ mice (red color). Data are presented as mean ± SEM (n = 8 mice per group). Statistically significant changes were assessed by using the Two-way ANOVA repeated measurement statistical test with Sidak-Holm posthoc test. (**C**) Sections of pERK immunostaining in DRGs of TRPM3-WT and TRPM3-KO mice at 24 h after oxaliplatin or vehicle injection. Scale bars, 50 μm. (**D**) The percentage of pERK-positive cells in the DRG sections was counted (n = 6 sections, from three independent DRG preparations). Data are expressed as the mean ± SEM. Statistically significant changes were assessed by using the Two-way ANOVA statistical test with Sidak-Holm posthoc test. Where *p<0.05; **p<0.01; ***p<0.001;

Increased activity of DRGs neurons is associated with higher levels of the phosphorylated form of the pronociceptive protein ERK (pERK), which has been observed in animal models of inflammatory and neuropathic pain, including oxaliplatin-induced neuropathy [13],[15]. We therefore compared pERK expression analysed via immunohistochemical stainings of DRG neurons isolated from wild type and TRPM3^-/-^ mice 24 hr after dosing with oxaliplatin (6mg/kg) or vehicle. We found that the number of pERK-positive cells was significantly increased in DRG cell bodies of oxaliplatin-injected wild type animals compared to vehicle controls animals. Importantly, oxaliplatin did not cause any increase in pERK expression in DRGs from TRPM3^-/-^ mice (Fig. 5C, D). Taken together, these data indicate that TRPM3 deficiency protects mice from oxaliplatin-induced mechanical allodynia and cold hypersensitivity, and prevents the oxaliplatin-induced upregulation of the pronociceptive pERK.

### 3.4. Effects of TRPM3 inhibition on the development of CIPNP

To investigate whether acute pharmacological inhibition of TRPM3 affects CIPNP, we tested the effect the TRPM3 antagonist isosakuranetin [25] on oxaliplatin-induced cold and mechanical hypersensitivity (Fig. 6A, B), using the opioid tramadol (5mg/kg) as a positive control [17]. In these experiments, the cold and mechanical sensitivity of wild type mice was tested at baseline and at day 6 after oxaliplatin treatment, confirming significant hypersensitivity similar to the experiments shown in Fig. 5. After the first testing on day 6, animals received a single i.p. injection with isosakuranetin, tramadol or vehicle, followed by repeat testing for cold and mechanical sensitivity. In comparison with vehicle, both isosakuranetin and tramadol resulted in a significant increase in mechanical threshold and decrease in the cold nociceptive response (Fig. 6). Taken together, these findings indicate that TRPM3 antagonism alleviates oxaliplatin-induced cold and mechanical hypersensitivity.

**Fig. 6:**
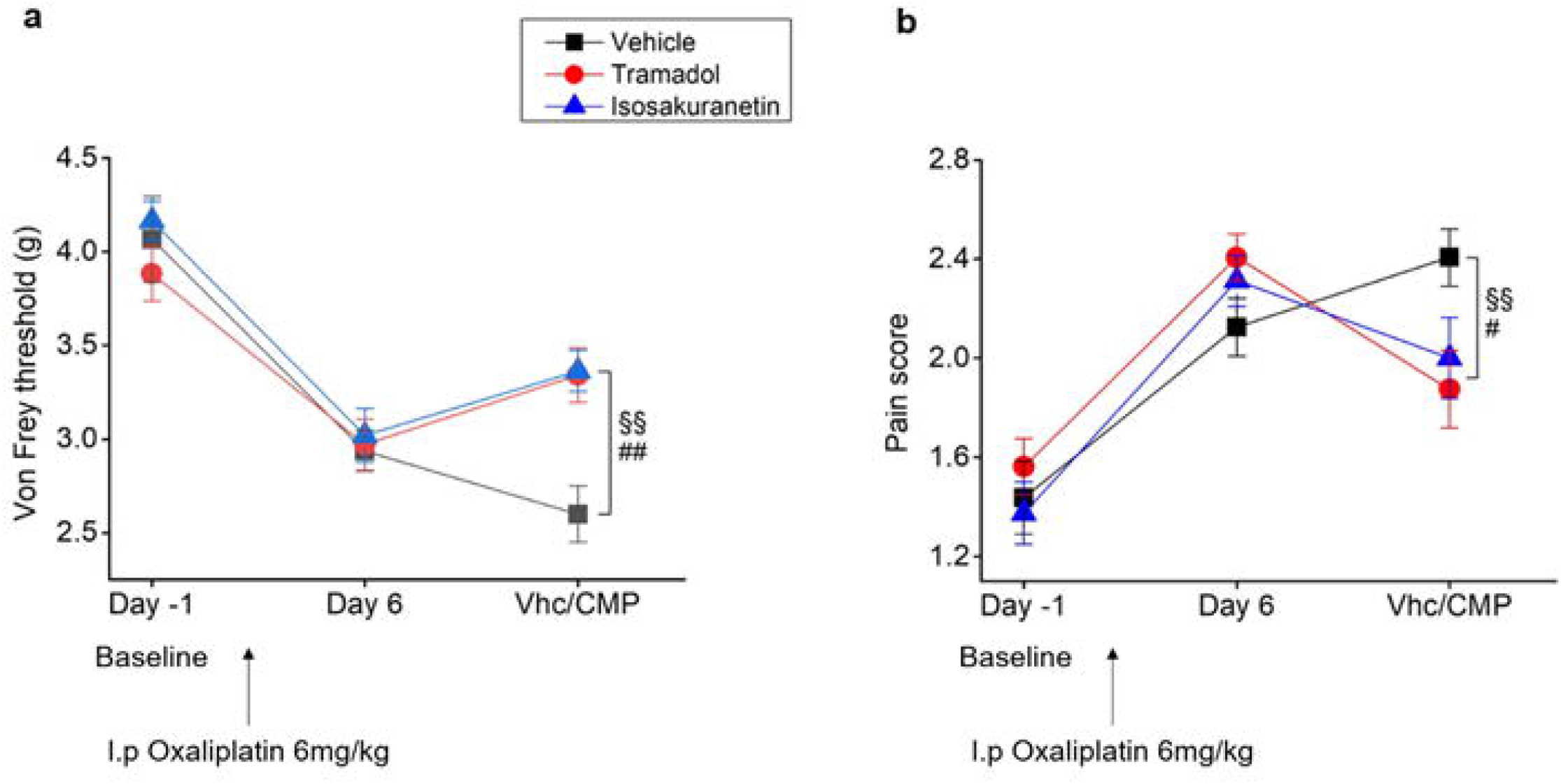
Pharmacological inhibition of TRPM3 reduces neuropathic pain induced by oxaliplatin. The effect is tested in the electronic von Frey assay and acetone spray test at day 6 post oxaliplatin treatment. (**A, B**) Effect of Isosakuranetin (2mg/kg), Tramadol as positive control (5mg/kg) and vehicle used in the intervention protocol on nociceptive mechanical threshold and escape behavior score in mice treated with oxaliplatin. Results are shown as mean ± SEM of n=8 mice per condition. Statistical analysis: Two-way ANOVA repeated measurements with Sidak-holm posthoc test. Where §§ vhc vs Tramadol (p<0.01); ## vhc vs Isosakuranetin (p<0.01); # vhc vs Isosakuranetin (p<0.05).

## 4. DISCUSSION

Oxaliplatin as a drug has been used for a long time in the treatment and management of cancer. Despite the effectiveness of oxaliplatin, the drug comes with substantial side effects, including hair loss, diarrhea, mouth sores, change in taste, dizziness, nosebleeds, and painful peripheral neuropathy. Several mechanisms have already been proposed in literature to explain the dose-limiting painful side effects often linked with the administration of chemotherapeutics agents such us oxaliplatin. For instance, oxaliplatin reportedly induces modifications of several intracellular signaling pathways [8], and alters activity of voltage-gated sodium channels and the sensory TRP channels TRPA1, TRPV1 and TRPM8 [19],[35]. Recently, another member of the TRP channel superfamily, TRPM3, has increasingly received attention as a potential key player in acute, inflammatory and neuropathic pain signaling and as a potential target for new analgesic treatments [26]. However, the contribution of TRPM3 to CIPNP remained unstudied. In this study, we present evidence that TRPM3 is critically involved in the development of acute oxaliplatin-induced neuropathic pain, and demonstrate that both genetic ablation and pharmacological inhibition of the channel significantly reverts oxaliplatin-induced mechanical hyperalgesia and cold hypersensitivity.

Oxaliplatin treatment was associated with increased TRPM3 activity, which we demonstrated in heterologous expression systems, in isolated sensory neurons as well as and in an *in vivo* chemogenic pain model. The mechanisms whereby oxaliplatin affects TRPM3 activity currently remain unknown. In contrast to earlier findings in inflammatory pain models [16], [33], [37], we did not find any evidence for increased TRPM3 expression at the mRNA level after oxaliplatin treatment. In contrast, we confirmed increased mRNA levels for the cold-activated TRPM8, similar to earlier studies [3],[6]. Thus, oxaliplatin enhances TRPM3 activity at the posttranscriptional level. Based on our experiments, we can also exclude a direct agonistic effect of oxaliplatin on TRPM3 channel function, suggesting that the enhanced channel activity is the consequence of increased protein expression at the plasma membrane or sensitized channel gating. In this respect, studies on other ion channels, including TRPV1 and TRPA1, point towards an important role for reactive oxygen species (ROS), which are generated following oxaliplatin-induced mitochondrial dysfunction [22] and can affect the activity of these channels through post-translational modifications to cysteine or proline residues. Further research is needed to test whether similar ROS modulation occurs at TRPM3. Oxaliplatin treatment in cancer patients frequently causes neuropathic pain, which encompasses cold hypersensitivity and mechanical allodynia. These symptoms develop within the first hours to days after dosing and can reproducibly be mimicked in rodent models. Here, we demonstrate a key role for TRPM3 in the development of these acute symptoms in mice. First, we show that the hypersensitivity towards cold and mechanical stimuli, which develops after oxaliplatin injection in wild type mice, is strongly suppressed in TRPM3^-/-^ mice. These findings establish that TRPM3 is required for the development of the hallmark symptoms of acute oxaliplatin-induced neuropathic pain. Moreover, we found that the oxaliplatin-induced upregulation of pERK in the cell bodies of DRG neurons, which is a biochemical marker of increased neuronal activation, is absent in TRPM3^-/-^ mice. In addition, we found that pharmacological inhibition of TRPM3 using isosakuranetin could effectively inhibit oxaliplatin-induced mechanical and cold hypersensitivity, with a similar efficacy as the opioid tramadol. Taken together, these findings reveal TRPM3 as a promising potential new target to treat painful hyperesthesia associated with oxaliplatin treatment.

Our observations that genetic ablation or pharmacological inhibition of TRPM3 have a strong effect on oxaliplatin-induced hypersensitivity to cold and mechanical stimuli may seem at odds with the known activation modalities of TRPM3 as a sensory ion channel. Indeed, TRPM3 is primarily known to be activated by and involved in the sensation of heat [31], [34] and specific chemical ligands; as such, cold temperatures reduce TRPM3 channel activity, and there is limited experimental evidence to link TRPM3 to the detection of mechanical stimuli. One possible explanation for our findings could be that the oxaliplatin-induced increased functionality of TRPM3, a depolarizing cation current, increases the excitability of the large set of TRPM3-expressing DRG neurons, including those involved in detecting cold and mechanical stimuli. In such a scenario, inhibition of TRPM3 function would revert the sensory neuronal hyperexcitability, thereby reducing the thermal and mechanical sensitivity to control levels. A similar role for TRPM3 in modulating nociceptor excitability has also been proposed in a mouse model of inflammatory hyperalgesia [25]. In those experiments, inhibition of TRPM3 in neurons innervating inflamed tissue reduced the responses to agonists of TRPA1 and TRPV1, which are broadly co-expressed with TRPM3 in nociceptor neurons.

In conclusion, we have demonstrated that the chemotherapeutic drug oxaliplatin evokes enhanced TRPM3 activity *in vitro* and *in vivo*, and is essential for the development of oxaliplatin-induced cold and mechanical hypersensitivity. TRPM3 thus represents a potential new target for the treatment of neuropathic pain in patients undergoing chemotherapy.

## 6. ACKNOWLEDGEMENT

We thank all members of the Laboratory of Endometrium, Endometriosis and Reproductive Medicine (LEERM), the members of the Laboratory of Ion Channel Research (LICR). Specific thanks to the members of the Laboratory of Neurobiology in particular professor Ludo van Den Bosch and Stijn Verschoren (KU Leuven) for their help during experiments and useful discussions during lab meetings.

## 7. FUNDING

We thank the Research Foundation-Flanders FWO, (G.0D1417N, G.084515N, G.0A6719N) and the Research Council of the KU Leuven (C14/18/106, C3/21/049) for funding the project of Joris Vriens.

## 8. CONFLICT OF INTEREST

The authors declare no conflict of interest

## 9. AUTHORS CONTRIBUTION

Conceptualization, T.V. and J.V.; methodology, V.D.A., S.P., R.V. and K.L.; validation, V.D.A. and S.P.; formal analysis, V.D.A.; investigation, V.D.A.; resources, T.V. and J.V; data curation, V.D.A.; writing - original draft, V.D.A.; writing - review and editing, V.D.A., T.V. and JV.; visualization, V.D.A and J.V.; supervision, T.V. and J.V; funding acquisition, T.V. and J.V. All authors have read and agreed to the published version of the manuscript.

## Notes

### Competing Interest Statement

The authors have declared no competing interest.

## BIBLIOGRAPHY

[1] Banach M, Juranek JK, Zygulska AL. Chemotherapy-induced neuropathies-a growing problem for patients and health care providers. Brain Behav 2017;7(1):e00558.

[2] Brzezinski K. Chemotherapy-induced polyneuropathy. Part I. Pathophysiology. Contemp Oncol (Pozn) 2012;16(1):72–78.

[3] Chukyo A, Chiba T, Kambe T, Yamamoto K, Kawakami K, Taguchi K, Abe K. Oxaliplatin-induced changes in expression of transient receptor potential channels in the dorsal root ganglion as a neuropathic mechanism for cold hypersensitivity. Neuropeptides 2018;67:95–101.

[4] Descoeur J, Pereira V, Pizzoccaro A, Francois A, Ling B, Maffre V, Couette B, Busserolles J, Courteix C, Noel J, Lazdunski M, Eschalier A, Authier N, Bourinet E. Oxaliplatin-induced cold hypersensitivity is due to remodelling of ion channel expression in nociceptors. EMBO Mol Med 2011;3(5):266–278.

[5] Fukuda Y, Li Y, Segal RA. A Mechanistic Understanding of Axon Degeneration in Chemotherapy-Induced Peripheral Neuropathy. Front Neurosci 2017;11:481.

[6] Gauchan P, Andoh T, Kato A, Kuraishi Y. Involvement of increased expression of transient receptor potential melastatin 8 in oxaliplatin-induced cold allodynia in mice. Neurosci Lett 2009;458(2):93–95.

[7] Held K, Kichko T, De Clercq K, Klaassen H, Van Bree R, Vanherck JC, Marchand A, Reeh PW, Chaltin P, Voets T, Vriens J. Activation of TRPM3 by a potent synthetic ligand reveals a role in peptide release. Proc Natl Acad Sci U S A 2015;112(11):E1363–1372.

[8] Joseph EK, Chen X, Bogen O, Levine JD. Oxaliplatin acts on IB4-positive nociceptors to induce an oxidative stress-dependent acute painful peripheral neuropathy. J Pain 2008;9(5):463–472.

[9] Kawashiri T, Egashira N, Kurobe K, Tsutsumi K, Yamashita Y, Ushio S, Yano T, Oishi R. L type Ca(2)+ channel blockers prevent oxaliplatin-induced cold hyperalgesia and TRPM8 overexpression in rats. Mol Pain 2012;8:7.

[10] Knowlton WM, Daniels RL, Palkar R, McCoy DD, McKemy DD. Pharmacological blockade of TRPM8 ion channels alters cold and cold pain responses in mice. PLoS One 2011;6(9):e25894.

[11] Krugel U, Straub I, Beckmann H, Schaefer M. Primidone inhibits TRPM3 and attenuates thermal nociception in vivo. Pain 2017;158(5):856–867.

[12] Levine JD, Alessandri-Haber N. TRP channels: targets for the relief of pain. Biochim Biophys Acta 2007;1772(8):989–1003.

[13] Liu DL, Lu N, Han WJ, Chen RG, Cong R, Xie RG, Zhang YF, Kong WW, Hu SJ, Luo C. Upregulation of Ih expressed in IB4-negative Adelta nociceptive DRG neurons contributes to mechanical hypersensitivity associated with cervical radiculopathic pain. Sci Rep 2015;5:16713.

[14] Martinov T, Mack M, Sykes A, Chatterjea D. Measuring changes in tactile sensitivity in the hind paw of mice using an electronic von Frey apparatus. J Vis Exp 2013(82):e51212.

[15] Maruta T, Nemoto T, Hidaka K, Koshida T, Shirasaka T, Yanagita T, Takeya R, Tsuneyoshi I. Upregulation of ERK phosphorylation in rat dorsal root ganglion neurons contributes to oxaliplatin-induced chronic neuropathic pain. PLoS ONE 2019;14(11): e0225586.

[16] Mulier M, Van Ranst N, Corthout N, Munck S, Vanden Berghe P, Vriens J, Voets T, Moilanen L. Upregulation of TRPM3 in nociceptors innervating inflamed tissue. Elife 2020;9.

[17] Naruge D, Nagashima F, Kawai K, Okano N, Kobayashi T, Furuse J. Tramadol/Acetaminophen Combination Tablets in Cancer Patients with Chemotherapy-Induced Peripheral Neuropathy: A Single-Arm Phase II Study. Palliat Med Rep 2020;1(1):25–31.

[18] Naziroglu M, Braidy N. Thermo-Sensitive TRP Channels: Novel Targets for Treating Chemotherapy-Induced Peripheral Pain. Front Physiol 2017;8:1040.

[19] Park SB, Lin CS, Krishnan AV, Goldstein D, Friedlander ML, Kiernan MC. Oxaliplatin-induced neurotoxicity: changes in axonal excitability precede development of neuropathy. Brain 2009;132(Pt 10):2712–2723.

[20] Raposo D, Morgado C, Pereira-Terra P, Tavares I. Nociceptive spinal cord neurons of laminae I-III exhibit oxidative stress damage during diabetic neuropathy which is prevented by early antioxidant treatment with epigallocatechin-gallate (EGCG). Brain Res Bull 2015;110:68–75.

[21] Sanchez JC, Munoz LV, Ehrlich BE. Modulating TRPV4 channels with paclitaxel and lithium. Cell Calcium 2020;91:102266.

[22] Santoro V, Jia R, Thompson H, Nijhuis A, Jeffery R, Kiakos K, Silver AR, Hartley JA, Hochhauser D. Role of Reactive Oxygen Species in the Abrogation of Oxaliplatin Activity by Cetuximab in Colorectal Cancer. J Natl Cancer Inst 2016;108(6):djv394.

[23] Smith EM, Pang H, Cirrincione C, Fleishman S, Paskett ED, Ahles T, Bressler LR, Fadul CE, Knox C, Le-Lindqwister N, Gilman PB, Shapiro CL, Alliance for Clinical Trials in O. Effect of duloxetine on pain, function, and quality of life among patients with chemotherapy-induced painful peripheral neuropathy: a randomized clinical trial. JAMA 2013;309(13):1359–1367.

[24] Soussain C, Ricard D, Fike JR, Mazeron JJ, Psimaras D, Delattre JY. CNS complications of radiotherapy and chemotherapy. Lancet 2009;374(9701):1639–1651.

[25] Straub I, Krugel U, Mohr F, Teichert J, Rizun O, Konrad M, Oberwinkler J, Schaefer M. Flavanones that selectively inhibit TRPM3 attenuate thermal nociception in vivo. Mol Pharmacol 2013;84(5):736–750.

[26] Su S, Yudin Y, Kim N, Tao YX, Rohacs T. TRPM3 Channels Play Roles in Heat Hypersensitivity and Spontaneous Pain after Nerve Injury. J Neurosci 2021;41(11):2457–2474.

[27] Taguchi K. [Role of Transient Receptor Potential Channels in Paclitaxel- and Oxaliplatin-induced Peripheral Neuropathy]. Yakugaku Zasshi 2016;136(2):287–296.

[28] Tatsushima Y, Egashira N, Kawashiri T, Mihara Y, Yano T, Mishima K, Oishi R. Involvement of substance P in peripheral neuropathy induced by paclitaxel but not oxaliplatin. J Pharmacol Exp Ther 2011;337(1):226–235.

[29] Van Helleputte L, Kater M, Cook DP, Eykens C, Rossaert E, Haeck W, Jaspers T, Geens N, Vanden Berghe P, Gysemans C, Mathieu C, Robberecht W, Van Damme P, Cavaletti G, Jarpe M, Van Den Bosch L. Inhibition of histone deacetylase 6 (HDAC6) protects against vincristine-induced peripheral neuropathies and inhibits tumor growth. Neurobiol Dis 2018;111:59–69.

[30] Van Hoeymissen E, Held K, Nogueira Freitas AC, Janssens A, Voets T, Vriens J. Gain of channel function and modified gating properties in TRPM3 mutants causing intellectual disability and epilepsy. Elife 2020;9.

[31] Vandewauw I, De Clercq K, Mulier M, Held K, Pinto S, Van Ranst N, Segal A, Voet T, Vennekens R, Zimmermann K, Vriens J, Voets T. A TRP channel trio mediates acute noxious heat sensing. Nature 2018;555(7698):662–666.

[32] Vangeel L, Benoit M, Miron Y, Miller PE, De Clercq K, Chaltin P, Verfaillie C, Vriens J, Voets T. Functional expression and pharmacological modulation of TRPM3 in human sensory neurons. Br J Pharmacol 2020;177(12):2683–2695.

[33] Vanneste M, Mulier M, Nogueira Freitas AC, Van Ranst N, Kerstens A, Voets T, Everaerts W. TRPM3 Is Expressed in Afferent Bladder Neurons and Is Upregulated during Bladder Inflammation. Int J Mol Sci 2021;23(1).

[34] Vriens J, Owsianik G, Hofmann T, Philipp SE, Stab J, Chen X, Benoit M, Xue F, Janssens A, Kerselaers S, Oberwinkler J, Vennekens R, Gudermann T, Nilius B, Voets T. TRPM3 is a nociceptor channel involved in the detection of noxious heat. Neuron 2011;70(3):482–494.

[35] Webster RG, Brain KL, Wilson RH, Grem JL, Vincent A. Oxaliplatin induces hyperexcitability at motor and autonomic neuromuscular junctions through effects on voltage-gated sodium channels. Br J Pharmacol 2005;146(7):1027–1039.

[36] Wickham R. Chemotherapy-induced peripheral neuropathy: a review and implications for oncology nursing practice. Clin J Oncol Nurs 2007;11(3):361–376.

[37] Zhao M, Liu L, Chen Z, Ding N, Wen J, Liu J, Ge N, Zhang X. Upregulation of transient receptor potential cation channel subfamily M member-3 in bladder afferents is involved in chronic pain in cyclophosphamide-induced cystitis. Pain 2022.

