## supplementary figures for "TRPM3 as novel target to alleviate acute oxaliplatin-induced peripheral neuropathic pain"

### Supplementary Material

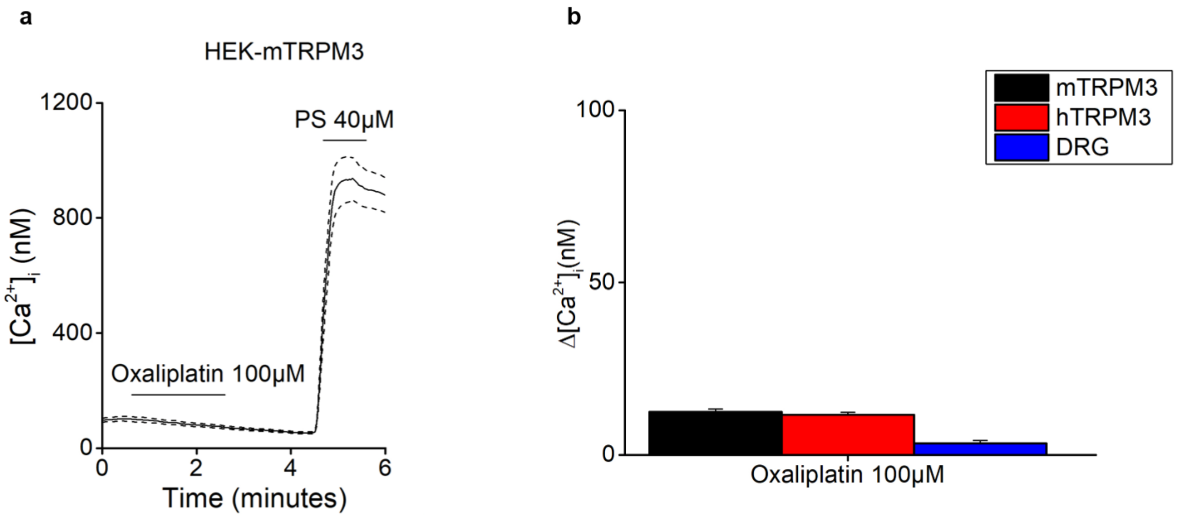


**Supplementary Fig. 1: The effect of acute addition of oxaliplatin on TRPM3 expressed in heterologous and homologous expression systems.** (**A**) Time course of intracellular calcium concentrations ([Ca$^{2+}$]i) (mean ± SEM) upon application of oxalipaltin (100 µM) and PS (40 µM) for HEK293-mTRPM3 cells. (**B**) Quantification of calcium responses upon direct application of oxaliplatin (100 µM) on HEK-mTRPM3 (black), HEK-hTRPM3 (red bar) and DRG neurons (blue bar).

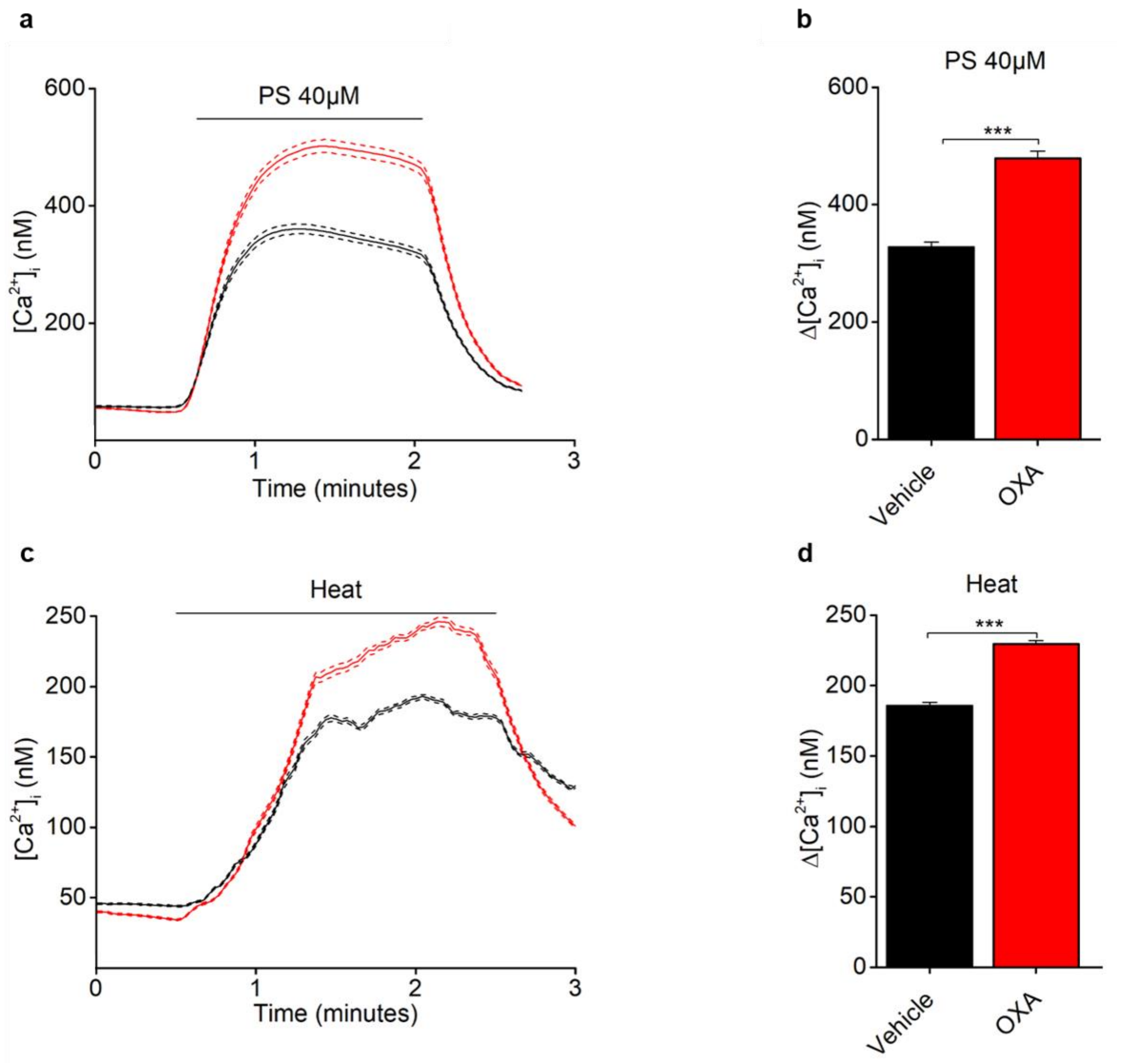


**Supplementary Fig. 2: Modulation of TRPM3 function in HEK293-hTRPM3 after oxaliplatin pretreatment**.

(**A** and **C**) Time course of intracellular calcium concentrations ([$Ca^{2+}$]i) (mean ± SEM) for HEK-hTRPM3 cells in response to PS (40µM) and heat (37°) after oxaliplatin (100µM; red bar) and vehicle (black bar) treatment. (**B** and **D**) Quantification of calcium responses for experiments as in panel A and C respectively. Where ***p=<0.001 (Mann-Whitney t test).

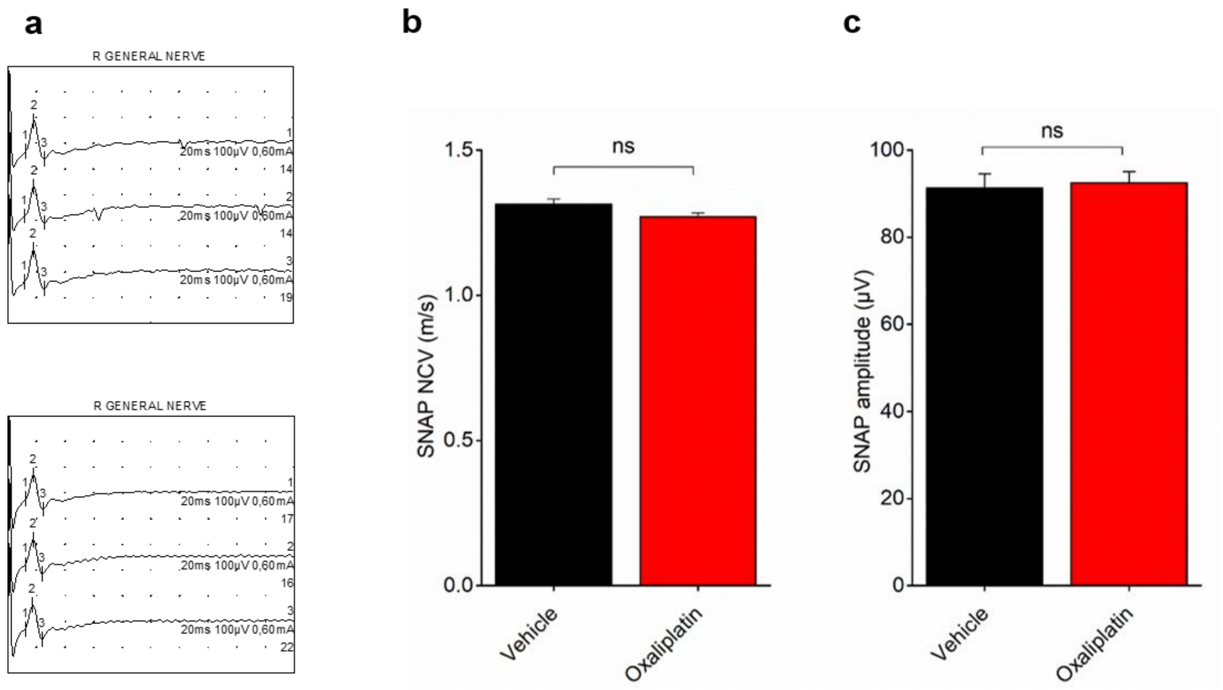


**Supplementary Fig. 3 Single administration of oxaliplatin does not affect the electrophysiological properties of caudal nerve conduction.**

(**A**) Typical sensory nerve conduction study graphs showing sensory nerve action potential traces and parameters such as the latency, the amplitude and the stimulus intensity. (**B, C**) Characterization of the nerve conduction velocity (NCV) and SNAP amplitude in the caudal nerve of mice treated with single i.p. injection of vehicle and Oxaliplatin (6mg/kg) for 24hr (n = 6 animals/ group).

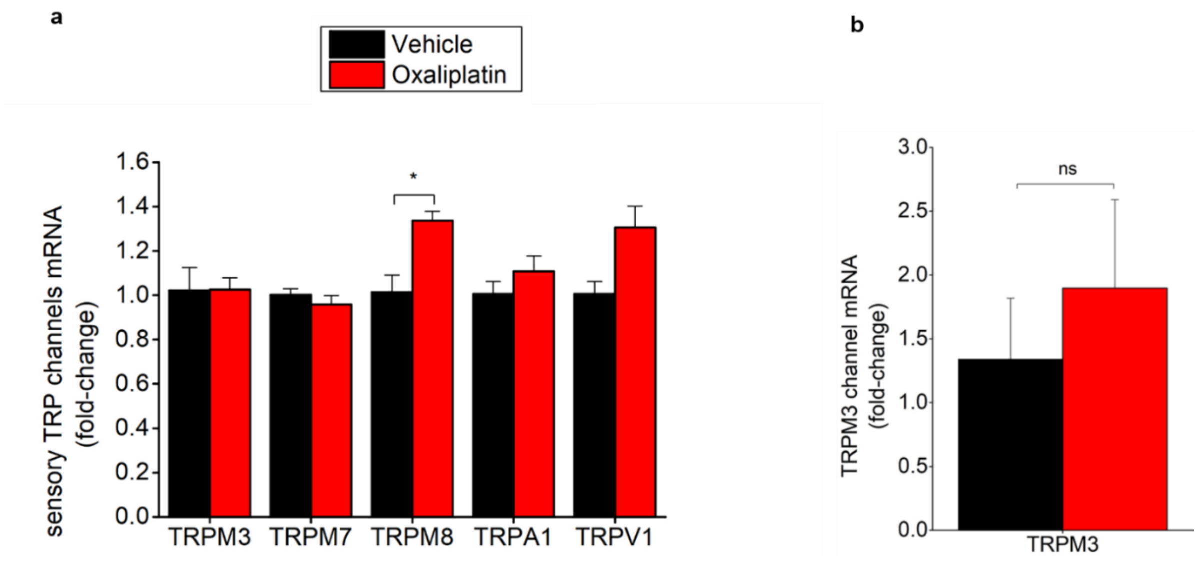


**Supplementary Fig. 4 Quantitative RT-PCR of TRP channels in primary DRG sensory neurons derived from vehicle and oxaliplatin treated mice after 24 hrs of treatment and HEK-mTRPM3.** mRNA levels were quantified to the geometric mean of housekeeping genes HPRT1 and PGK-1. Data are presented as mean ± SEM. Statistically significant changes in mRNA expression were assessed using the Mann-Whitney test. Where *p<0.05 (**A**) Quantitative RT-PCR showing the mRNA expression of the indicated TRP channels in isolated DRG treated with vehicle (black bars) and oxaliplatin (red bars)**.** (**B**) Quantitative RT-PCR showing the mRNA expression of TRPM3 in HEK-mTRPM3 cells treated for 24 hrs with vehicle (black) and oxaliplatin (red bars) (n = 3 independent experiments).
